# Melanism patches up the defective cuticular morphological traits through promoting the up-regulation of cuticular protein-coding genes in *Bombyx mori*

**DOI:** 10.1101/155002

**Authors:** Liang Qiao, Ri-xin Wang, You-jin Hao, Hai Hu, Gao Xiong, Song-zhen He, Jiang-bo Song, Kun-peng Lu, Ya-qun Xin, James Mallet, Bin Chen, Fang-yin Dai

**Author notes:** **Corresponding Authors: Liang Qiao**, Chongqing Key Laboratory of Vector Insects, Institute of Entomology and Molecular Biology, College of Life Sciences, Chongqing Normal University, No. 37 Daxuecheng Road, Chongqing, 401331, China, **Fang-yin Dai**, State Key Laboratory of Silkworm Genome Biology, Key Laboratory for Sericulture Functional Genomics and Biotechnology of Agricultural Ministry, Southwest University, No. 2 Tiansheng Road, Chongqing, China.

## Abstract

Melanin and cuticular proteins are important cuticle components in insect. Cuticle defects caused by mutations in cuticular protein-encoding genes can hinder melanin deposition. However, the effects of melanin variation on cuticular protein-encoding genes and the corresponding morphological traits associated with these genes are remain largely unknown. Using *Bombyx mori* as a model, we showed that the melanism levels during larval cuticle pigmentation correlated positively with the expression of cuticular protein-encoding genes. This correlation stemmed from the simultaneous induction of these genes by the melanin precursors. More importantly, the effect of the melanism background on the cuticles induced the up-regulation of other functionally redundant cuticular protein-encoding genes to rescue the morphological and adaptive defects caused by the dysfunction of some mutated cuticular proteins, and the restorative ability increased with increasing melanism levels, which gives a novel evidence that melanism enhances insect adaptability. These findings deepen our understanding of the interactions among cuticle components, as well as their importance in the stabilizing of the normal morphology and function of the cuticle.

## Introduction

The prerequisite for the benefits of melanism to insect is not only the integrity of the melanin biosynthesis and regulatory pathway (Wilson *et al*. 2001; Liu *et al*. 2015; Mallet and Hoekstra 2016), but also the normal presence of the platform it relied on (Wittkopp *et al*. 2003; Wittkopp and Beldade 2009; Andersen 2010; Moussian 2010; Van Belleghem *et al*. 2017). For the insects, the most important fundamental platform for the color pattern drawing is the cuticle (Hopkins and Kramer 1992; Andersen 2010; Moussian 2010). During shaping the cuticle, the maintenance and stability of the cuticle features depends on normal functional cuticular proteins and the interactions with other components (such as chitin) (Hopkins and Kramer 1992; Guan *et al*. 2006; Suderman *et al*. 2006; Andersen 2010; Moussian 2010; Chaudhari *et al*. 2011; Noh *et al*. 2016). Due to the crucial roles of cuticular proteins on cuticle development, when their coding genes are loss of function, the abnormal or defective cuticle will likely affect the deposition and attachment of melanin, which is not conducive to the performance of color pattern (Kanekatsu *et al*. 1988; Oota and Kanekatsu 1993; Arakane *et al*. 2012; Jasrapuria *et al*. 2012; Wang 2013; Noh *et al*. 2015). Yet little is known about the corresponding response mode of cuticular protein-encoding genes via the alteration of the melanin biosynthesis or regulatory pathway.

Recently, several high throughput expression surveys showed that the abundance of cuticular protein-encoding genes in different colored integuments varied in some insect species, especially with evidently up-regulated in the melanic regions (Futahashi *et al*. 2012; He *et al*. 2016; Wu *et al*. 2016; Tajiri 2017), and some of those shared very similar expression patterns and functions (Nakato *et al*. 1994; Nakato *et al*. 1997; Shofuda *et al*. 1999; Okamoto *et al*. 2008; Liang *et al*. 2010; Tang *et al.* 2010; Qiao *et al*. 2014).These studies implied that there are probably some relationships between the promotion of melanism and the expression of cuticular protein-encoding genes. Prior to this study, further exploring and understanding of the potential relationships were unclear. Additionally, when melanism-promoting instructions and defective cuticle proteins occur simultaneously, what are the effects of their relationships on the presence of the corresponding morphological traits?

In the Lepidoptera model, *Bombyx mori*, an intriguing phenomenon has been reported that a larval melanic mutant, *Striped* (*p^S^*, 2-0.0) is able to rehabilitate the malformed body shape, as well as the adaptability defects of the *stony* mutant (*st*, 8-0.0) (Xiang 1995; Banno *et al*. 2005). A recent study clarified that a transcription factor, *Apontic-like*, which boosts the expression of melanin synthesis genes in epidermal cells, is responsible for the *p^S^* mutant (Yoda *et al*. 2014). Besides this, there are also multiple alleles with different melanism degrees at the *p* locus, including *p^B^, p^M^, etc* (Xiang 1995; Banno *et al*. 2005; Yoda *et al*. 2014). And the *stony* mutant (*st*, 8-0.0) which precisely caused by the deletion of a RR1-type larval cuticular protein-encoding gene, *BmLcp17* (or *BmorCPR2*) showed hard and tight touch feeling, uncoordinated ratios of the length of the internodes and the intersegment folds (I/IF), bulges at intersegment folds, and severely defective locomotion and behavioral activities in the larval stage (Qiao *et al*. 2014). In addition, the similarity of the gene expression patterns and functional characteristics among some members of the RR1-type larval cuticular protein-encoding gene family, such as *BmLcp18, BmLcp22, BmLcp30* also suggest that they may play very similar roles as *BmLcp17* in shaping the larvae cuticle (Nakato *et al*. 1994; Nakato *et al*. 1997; Shofuda *et al*. 1999; Okamoto *et al*. 2008; Liang *et al*. 2010; Tang *et al*. 2010; Qiao *et al*. 2014). These dispersed findings are linked through the epistasis of *p^S^* to *stony*, then provide the breakthrough and the exceptional genetic resources for exploring the interactions between melanin and cuticular protein-encoding genes.

Here we illustrated that the transcripts levels of four cuticular protein-encoding genes were positively correlated with the melanism degree of larval cuticle, which were due to the simultaneous induction these genes by the intracellular melanin precursors. Moreover, by importing melanism-promoting instruction into the *stony* mutant, the cuticle deficiency rescued through functionally redundant compensation by some other up-regulated cuticular protein-encoding genes, which a new evidence that melanism as a beneficial trait. These findings deepen our understanding of the interactions among genetic factors which shape morphological features in lepidopteran, and emphasize the ecological and evolutionary significance of these mutual interactions.

## Materials and Methods

### Silkworm strains

Wild-type strains Dazao (+^*p*^) and melanic mutant strains *p^M^, p^S^* and *p^B^* (Xiang 1995; Banno *et al*. 2005; Yoda *et al*. 2014) were analyzed in this study. The darkness of pigment was measured as mean OD value using Image J (https://imagej.nih.gov/ij/). In terms of melanism degree, the body color of an individual that is homozygous at one of the aforementioned melanic loci is darker than that of a heterozygous individual (Xiang 1995; Banno *et al*. 2005). The albinism mutant *albino* (*al*) (Banno *et al*. 2005; Fujii *et al*. 2013), non-diapause wild-type strain N4 (used for melanin inhibition treatment) and *BmLcp17* deletion strain Dazao-*stony* (near isogeneic line of Dazao, which have been backcrossed with Dazao over 26 generations) were supplied by the Silkworm Gene Bank in Southwest University. The N4 strain and *al* mutant were fed with artificial diet at 28°C, and the others were fed fresh mulberry leaves under a 12□h/12□h light/dark photoperiod at 24□°C.

### Chemicals and cell lines

L-dopa (D9628), dopamine (H8502), tetrahydrofolic acid (BH_4_) (T3125) and 2,4-Diamino-6-hydroxypyrimidine (DAHP) (D19206) were purchased from Sigma-Aldrich (St. Louis, MO, USA). *BmNs* cell line was kept in our laboratory.

### Mating combinations and progeny phenotypes identification

The *p^S^* and *p^M^* strains were crossed with the Dazao-*stony* strain to generate the F_1_ generation, respectively. The F_2_ generation were produced by an F_1_ self-cross, and individuals of day 5 of the fifth-instar were collected to further use. The *p^B^* strain was crossed with the Dazao-*stony* strain to generate F_1_ progeny, which mated with the Dazao-*stony* strain to generate the BC_1_ generation, and fed them until at day 5 of the fifth-instar. Firstly, individuals of the F_2_ or BC_1_ generations were separated by their cuticle pigmentation. Subsequently, their phenotypes were further classified by morphological characteristics, touch feeling, and the ratios between the number of internodes and intersegmental folds in the second, third, and fourth abdominal segments based on an earlier method (Qiao *et al*. 2014).

### Genotyping

Because the *p^S^, p^M^* and *p^B^* mutations are alleles at the *p* locus, they should be located in proximity each other on the chromosome 2 (Xiang 1995; Banno *et al*. 2005; Yoda *et al*. 2014). Based on the reported sequence of the gene corresponding to the *p^S^* allele, a polymerase chain reaction (PCR)-based molecular marker within of the genomic region of the *Apontic-like* was designed. PCR screening were performed in the *p^X^* (*X*=*M, S* and *B*) and Dazao-*stony* to obtain the polymorphism molecular marker for *p* locus. Similarly, molecular marker was also designed within genomic region of the *BmLcp17* for polymorphism screening of the *stony* locus. The primers used in this study are listed in Table S1.

### Association analysis of gene expression, phenotype and genotype

Day 5 fifth-instar larvae of the Dazao or Dazao-*stony* strains were selected for cuticle dissection. The cuticles of the semi-lunar marking region and the non-melanic portion between the paired semi-lunar marking were finely dissected. Gene expression levels of *BmLcp17, BmLcp18, BmLcp22* and *BmLcp30* in these regions were compared. Gene expression patterns were determined for the dorsal epidermis of abdominal segments (from semi-lunar marking to star marking) from day 5 of fifth instar larvae (Dazao, *p^S^/+, p^M^/+, p^B^/+*). Day 1 of second instar larvae of the *al* and Dazao strains were also investigated. The same regions of dorsal epidermis regions were collected from F_2_-generation individuals with the *p^S^/p^S^, st/st*, and *p^S^/+, st/st* genotypes, as well as the *p^M^/p^M^, st/st* and *p^M^/+, st/st* genotypes for gene expression analyses. For all genotyped individuals, the ratios of the intersegment length to the intersegment fold were also analysed using image J.

### Melanin precursors-promoting and inhibition treatments

The preparation and concentration of L-dopa and dopamine solutions were slightly modified according to the description by Futahashi (Futahashi and Fujiwara 2005), and filtered with a 0.22 μm membranes before use. Cells were washed three times with Grace medium without melanin precursors to remove metabolites and other contaminants on the cell surfaces. Medium (0.8 mL) containing L-dopa or dopamine was added separately into each well, and the medium without melanin precursors was used as the control. Culture plates were sealed with tape, wrapped with foil and incubated at 28°C for 24 h for a gene expression analysis. For BH_4_ feeding assays, the 30mM working solution was prepared by dissolving tetrahydrofolic acid into ddH_2_O, and spreading it on an artificial diet for *al* mutants. The control was ddH2O. Phenotype were observed and recorded from the second instar stage, and expression of cuticular protein-encoding genes was analysed. For the melanism-inhibition experiments, the wild-type strain N4 was used. Newly-hatched larvae were divided into treatment and control groups. Individuals in the treatment group fed with DAHP dissolved in 0.1M NaOH (15g/L), and individuals in the control group fed 0.1M with NaOH. Phenotypes observation and gene expressions detection were performed on day 1 of the second-instar larvae.

### Quantitative RT–PCR

Total RNA extraction, reverse transcription and Quantitative reverse transcription-PCR (qRT-PCR) conducted as described previously (Qiao *et al*. 2014). Three biological replicates were prepared for each condition, and *BmRPL3* was used as the internal control. Primers are listed in Table S1.

## Results

### Entirely distinct expression patterns of cuticular protein-encoding genes and the cuticle appearance between melanic and non-melanic regions

Expression patterns of *BmLcp17, BmLcp18, BmLcp22* and *BmLcp30* in melanic or non-melanic epidermal regions are shown in Figure 1. These results, together with earlier studies (Futahashi *et al*. 2012; He *et al*. 2016; Wu *et al*. 2016), revealed that the gene expressions were significantly higher in melanic parts of the cuticle than in non-melanic parts. Moreover, micro protrusions were more intensive in the melanic regions than in the non-melanic regions, and accompanied by a higher amounts of chitin (another important cuticle component (Hopkins and Kramer 1992; Moussian *et al*. 2006; Andersen 2010; Moussian 2010; Chaudhari *et al*. 2011; Qiao *et al*. 2014), and the reduction chitin content was reported to impede cuticle melanism (Moussian *et al*. 2005)) (Figure S1A). These results showed that, regardless of the different genetic basis of the melanic mutants or the melanic markings in the non-melanic strains, excessive accumulation of melanin in the cuticle (accompany by the changes in the surface structure and the chitin content of the melanic cuticle) was closely related to the high expression levels of the cuticular protein-encoding genes.

**Figure 1.**
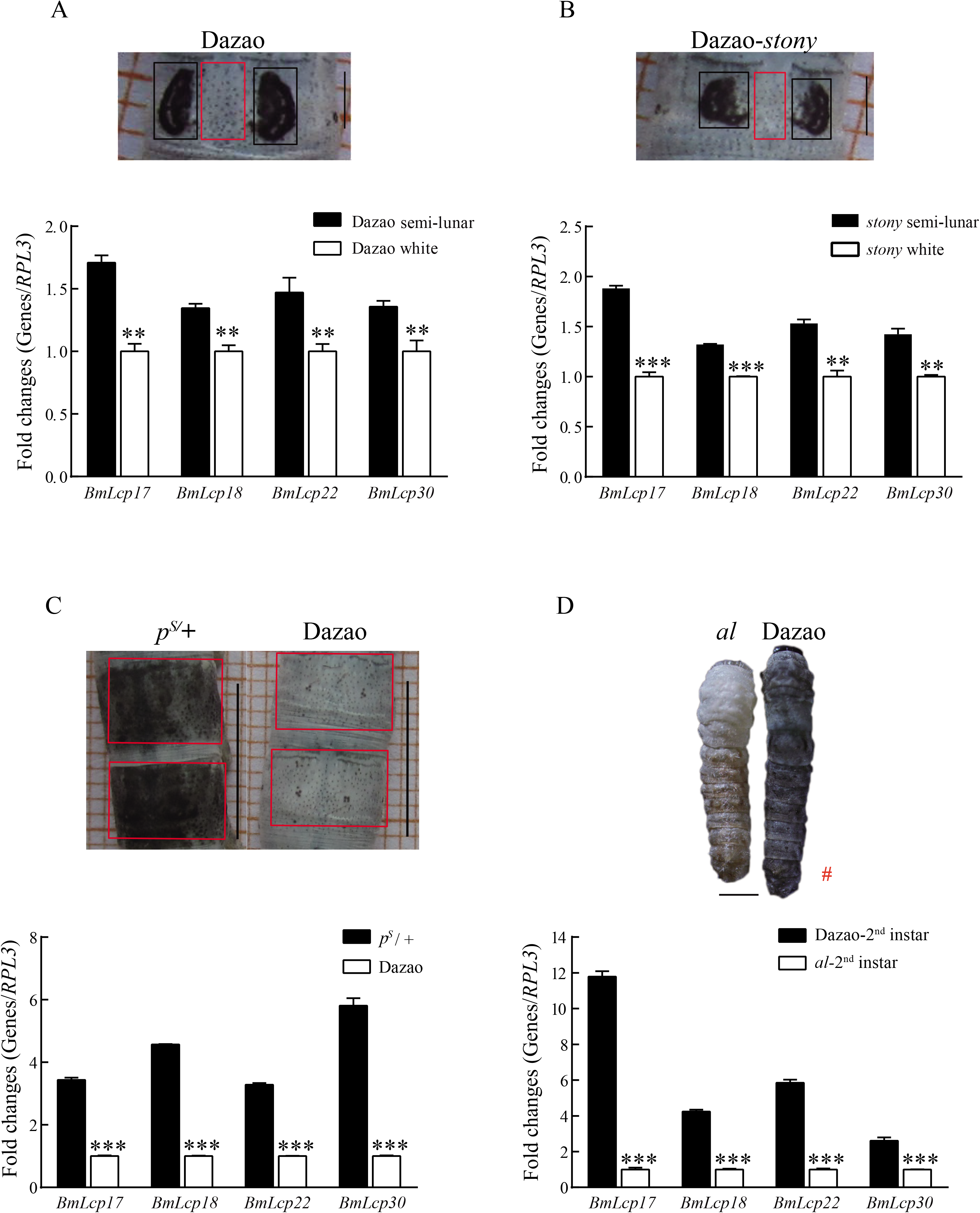
Expression of four larval cuticular protein-encoding genes in melanic and non-melanic integuments. A and B represent the gene expressions between the semi-lunar marking (black box) and the non-melanic region (between the semi-lunar marking, red box) in Dazao or Dazao-*stony*, respectively. Scale bar: 2 mm. C. Comparative analysis of gene expressions in the dorsal side of abdominal segments (from the third to the fourth segment, red box) between the *p^S^* and Dazao strains. Scale bar: 1 cm. D. Comparison of gene expressions between the second-instar *al* mutant and the Dazao strain (melanic). The red hashtag symbol indicates the Fig. 1 D we are showing is cited from the previous study of our lab group (Min *et al*. 2016) with modification. Scale bar: 2 mm. *t*-test, n=3; * *p* < 0.05; ** *p* < 0.01; *** *p* < 0.001.

### Expression level of larval cuticular protein-encoding genes positively correlated with the degree of cuticle melanism

To obtain further insights into the relationships between the melanin accumulation and the expression of cuticular protein-encoding genes, we investigated gene expression patterns using four *p* locus alleles (Dazao (+^*p*^), *p^M^, p^S^* and *p^B^*), which showed gradually increasing melanin accumulation (Figure 2A). Gene expression levels were gradually and significantly up-regulated with the increase in the degree of melanism in cuticle (Figure 2B). These results further showed that the expression levels correlated positively with the degree of melanism. Thus, the quantities of cuticular proteins with similar or redundant functions was increased greatly by the increasing the degree of melanism.

**Figure 2.**
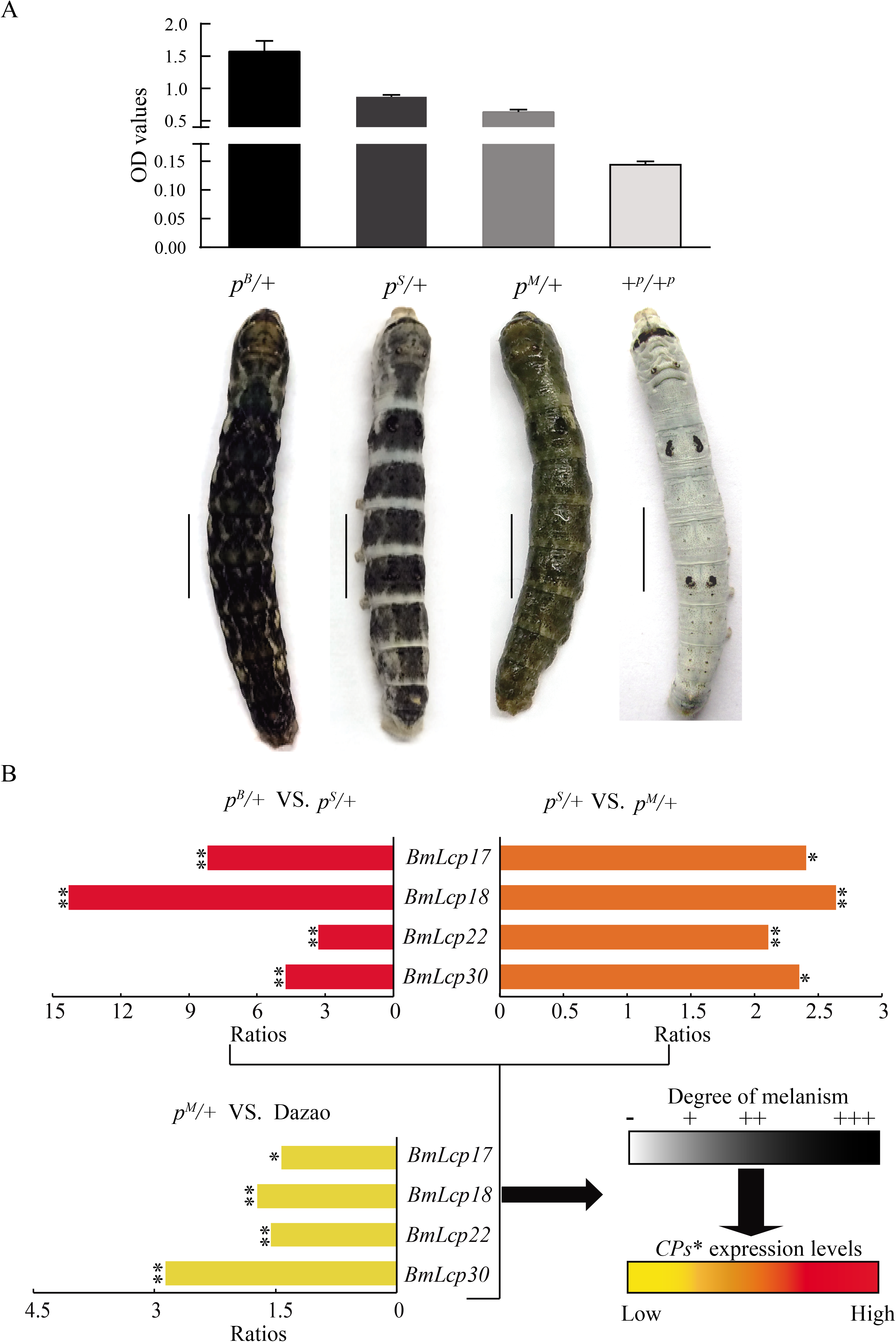
Expression of cuticular protein-encoding genes in integuments showing different degree of melanism. A and B display comparisons of degree of melanism and cuticular protein-encoding gene expression levels among four strains with mutant alleles at the *p* locus. Scale bar: 1 cm. Ratios represent the ratios of gene expression levels between two strains. Symbols (−, +, ++, and +++) represent the increment of the degree of melanism. Star represents the melanin-associated cuticular protein-encoding genes. *t*-test, n=3; * *p* < 0.05; ** *p* < 0.01.

### Typical stony phenotyped individuals masked after the introduction of melanic loci into the BmLcp17 defection strain

We assessed the effects of modulating the melanic background on the phenotypic defects caused by the deletion of *BmLcp17*. After mating the *p^B^* and *stony* parental, the percentage of BC_1_ individuals with melanism and the normal body shape in the backcrossed population of the *p^B^*×*stony* cross was 52 % (290/553; and theoretically, it was 25 %), yet individuals with the melanism cuticle and *stony*-type body shape were not found (theoretically should be almost equivalent to the number of individuals with melanism cuticle and normal body shape) (Figure 3). In the cross of *p^S^*×*stony*, ∼10.8% (36/331) of F_2_ progeny had an lighter melanism body color and smaller body size (Figure 3). These individuals exhibited a little hard and tight touched body, but the body was much softer than that of the *stony* mutant. Their intersegment folds bulged slightly, and the length were significantly shorter than that of the internodes; accordingly, their phenotypes slightly resembled the morphological features of the *stony* mutant (Figure 3 and 4A). Even so, We did not find individuals with the typical *stony*-type body shape and also defective adaptability under the melanism background (theoretical ratio is 3/16). (Figure 3). Similarly, in the *p^M^*×*stony* corss, only approximately ∼11.7% (51/437) of the individuals of the F_2_ population were very lightly melanic, but they exhibited obviously unusual morphological features (Figure 3, Figure 4A). When compared with the ∼10.8% F_2_ individuals (aforementioned) from the *p^S^*×*stony* cross, the touch feelings of individuals from the *p^M^*×*stony* cross were tighter and firmer, and the intersegment folds bulged more obviously and had a higher proportion among the segments, meaning that their body features were closer to the phenotype of the *stony* mutant (Figure 3 and 4A) (Qiao *et al*. 2014). Nevertheless, melanic individuals showing the typical *stony*-type body features and defective adaptability still did not appear (theoretical ratio is 3/16) in the progeny from the *p^M^* × *stony* cross. Therefore, induction of the melanic mutation into individuals with a defective cuticular protein-encoding gene could mask the adverse phenotypes, and the masking effect was more remarkable with the increasing degree of melanism.

**Figure 3.**
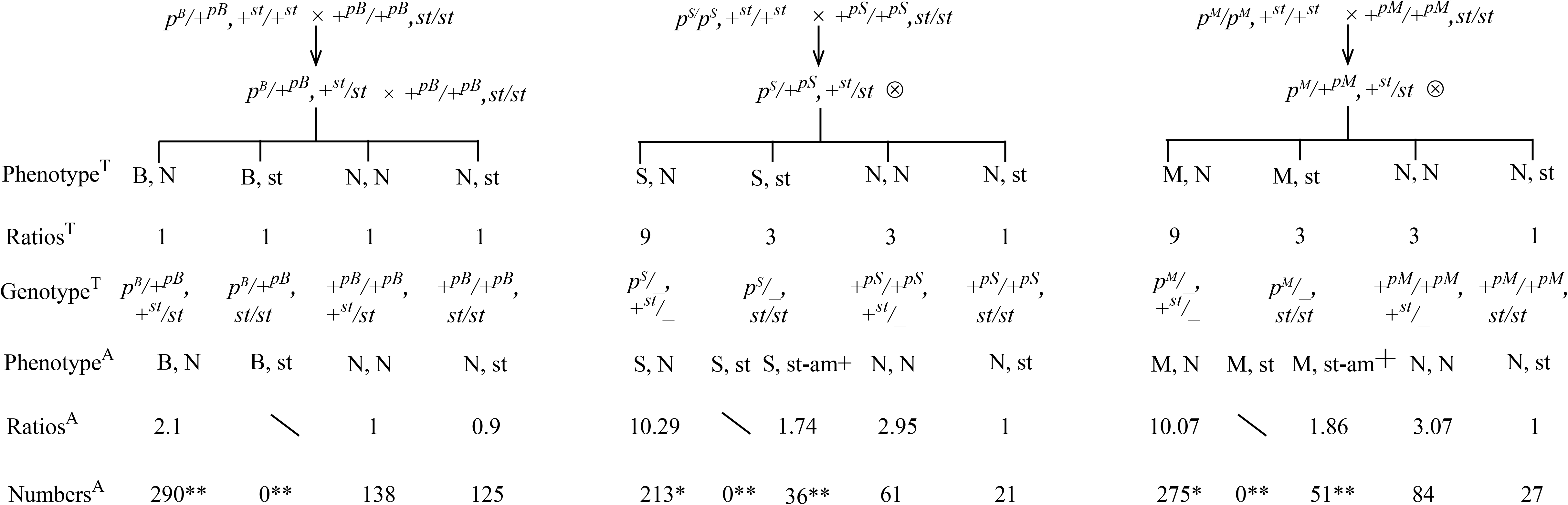
Segregation patterns of the phenotypes of progenies from different crosses of melanic mutant strains and the *stony* strain. In the segregated progenies, the first item in the list (B,N), (B,st), (S,N), (S,st), (M,N), (M,st), (N,N) and (N,st) represents *p^B^-, p^S^ -p^M^*- and Normal color patterns, respectively. The second item indicates body shape features marked with non-*stony* type (N) and the *stony* type (st). It is noteworthy that in (S, st-am+) and (M, st-am+), the second item represents the ambiguous *stony* body shape. The size of “+” symbol represents the corresponding degree of the ambiguous *stony* body shape. Superscript “T”s represent theoretical values. erscript “A”s represent actual values. Backslashes indicates that a value was not obtained. Chi-squared test, * *p* < 0.05; ** *p* < 0.01.

**Figure 4.**
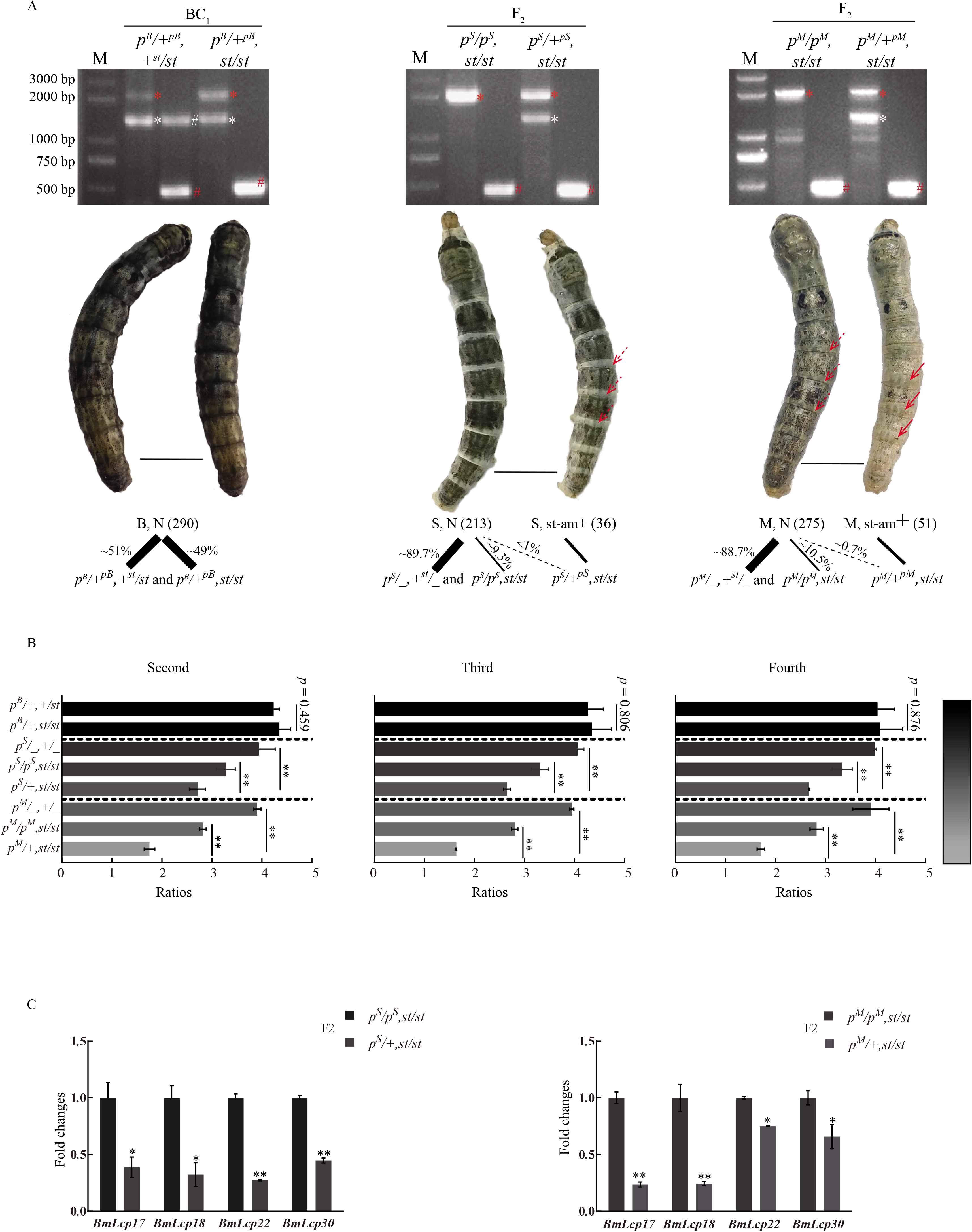
Association analysis of the genotypes, phenotypes and gene expression levels in segregated progenies from different crosses. A. Correlation analysis between the genotypes and phenotypes in self-crossed or backcrossed progenies. Scale bar: 1 cm. White and red stars represent polymorphic bands at the +^*p*^ and *p^X^* loci (*X* = *B, S* or *M*), respectively. Red and white hash-tag represent polymorphic bands at the *st* and +^*st*^ locus, respectively. Solid and dotted red arrows indicate the relative degree of bulging (solid > dotted), respectively. Slashes show the genotypes within one phenotypic category. The thickness of the slash represents the proportion of the corresponding genotypes. Dotted backslashes indicate that the number of individuals with the corresponding genotype is quite low. B. Ratios of the length of internodes and intersegmental folds in the second, third, and fourth abdominal segments of individuals with different genotypes (showing melanic body color) in the self-crossed or backcrossed progenies. n≥3, *t*-test, ** *p* < 0.01. C. Gene expression analysis of cuticular protein-encoding genes in homogeneous and heterogeneous individuals at the *p* locus from self-crossed progenies of *p^S^* × *stony*, and *p^M^* × *stony* under the condition that the cuticle was melanic and the genotype was homozygous recessive at *st* locus. n = 3, *t* – test, * *p* < 0.05; ** *p* ≤ 0.01.

### Other larval cuticular protein-encoding genes up-regulated evidently in the melanic and non-*stony* phenotypic, but with mutated BmLcp17 genotypic offspring

Using the molecular markers (closely linked to the *p* and *st* loci, respectively), we further genotyped the progenies with melanic color pattern and non-*stony* (including ambiguous *stony*-like) body shape. The results revealed that approximately 49% of the individuals showing a melanic color and normal body shape in the BC_1_ population from the *p^B^*×*stony* cross was the *p^B^*/+^*pB*^, *st/st* genotype (Figure 4A). The ratios of the length of the internodes and the intersegment folds (I/IF) were ∼4, which is very similar to that in *p^B^*/+^*pB*^, +^*st*^/*st* individuals, and also no significant differences as that in the wild-type individuals (Figure 4B) (Qiao *et al*. 2014). In the F_2_ generation from the *p^S^*×*stony* cross, approximately 9.3% of the individuals with the *p^S^*/*p^S^, st*/*st* genotype and very few individuals (∼1%) with the genotype *p^S^*/+^*pS*^, *st/st* exhibited a melanic color and the normal body shape (Figure 4A). For *p^S^*/*p^S^, st/st* genotyped individuals, the I/IF value was approximately 3.3 (Figure 4B). Although the I/IF value was lower than that in *p^S^*/_, +^*st*^/_ individuals (approximately 4, Figure 4B), it was much higher than that in the *stony* mutant (approximately 1.6 (Qiao *et al*. 2014)). Despite the slightly smaller body size of the *p^S^*/*p^S^, st/st* individuals, their body shape was essentially normal (Figure 4A). The genotypes of those lightly melanic individuals, whose body shape was slightly like that of the *stony* phenotype (mentioned in Result 3), were all the *p^S^*/+^*pS*^, *st*/*st*, and the I/IF value of these individuals was approximately 2.7 (Figures 4A and 4B). In progeny of the *p^M^*×*stony* cross, ∼10.5% of the individuals with the *p^M^/p^M^, st/st* genotype and ∼0.7 % of the individuals with the *p^M^*/+^*pM*^, *st/st* genotype showed a melanic color and subtle *stony* features (just very slight bulges) (Figure 4A). The body size of the *p^M^*/*p^M^, st/st* individuals were smaller than those of the *p^M^*/_, +^*st*^/_ individuals. They exhibited a certain sense of hardness, and the I/IF ratio was approximately 2.8, which is in good agreement with their phenotypes (Figure 4B). Additionally, individuals showing very slight melanism and morphological features that were more similar to that of the *stony* mutant (mentioned in Result 3) were all the *p^M^*/+^*pM*^, *st/st* genotype, and their I/IF values were approximately 1.8, which is closer to that of the *stony* mutant (Figures 4A and 4B). In addition, the expression of cuticular protein-encoding genes in *p^S^/p^S^, st/st* individuals were significantly higher than that in *p*^*S*^ /+^*pS*^, *st/st* individuals (Figure. 4C); a similar result was also obtained from the *p^M^*/*p^M^, st/st* and *p^M^*/+^*pM*^, *st/st* individuals (Figure 4C). Taken together, these results revealed that more cuticular proteins with similar functions were accumulated in the cuticle of melanic homologous individuals at the *p* locus. Based on the comprehensive and association analysis, we infer that melanic background effectively drove the expressions of cuticular protein-encoding genes with similar expression patterns and redundant functions, which compensated for the morphological and adaptability defects caused by the dysfunctional *BmLcp17* gene; and the law of compensatory abilities was *p^B^*/+^*pB*^, *st/st* > *p^S^*/*p^S^, st/st* > *p^M^/p^M^, st/st* **≮** *p^S^*/+^*pS*^, *st/st* > *p^M^*/+^*pM*^, *st/st*, which corresponds well with the gradual weakening of the degree of melanism (Figures 4 and Figure S3).

### Content variations of melanin precursors affect the transcript amount of the cuticular protein-encoding genes

Due to the crucial material basis for the cuticle melanism (no matter what kind of genetic basis caused by) is the extensive accumulation of melanin precursors in the epidermal cells; thus, the essence that melanism tend to increase the expression of some cuticular protein-encoding genes should be driven by the relations between the accumulation of melanin precursors and the transcripts amount of the cuticular protein-encoding genes. The basal expressions of four cuticular protein-encoding genes were detected in *BmNs* cells (Figure S5) and indicated that regulatory pathways controlling the expression of cuticular protein-coding genes in this cell line. Thus, this cell line was used to examine the effect of melanin precursors on gene expressions. After incubating *BmNs* cells with melanin precursors, the expression of cuticular protein-encoding genes was significantly higher in cells treated either by L-dopa or dopamine, compare with that in the control group (Figure 5 top (left)). Second-instar *al* mutant are characterized by albinism and a porous cuticle due to a mutation in sepiapterin reductase, which leads to the insufficient synthesis of the co-factor BH_4_ during the synthesis of melanin precursors (Banno *et al*. 2005; Fujii *et al*. 2013). When treated with BH_4_, these mutants had normal melanic body color (Lab unpublished contribution) (Fujii *et al*. 2013), and gene expressions were obviously higher than that in the control group (Figure 5 top (right)). These results suggested that the expression of cuticular protein-encoding genes was induced by increasing amounts of melanin precursors. We investigated the effects of DAHP, an inhibitor of guanylate cyclase hydrolase (GTPCHI), an important rate-limiting enzyme in the synthesis of BH_4_ (Hamadate *et al*. 2008). Inhibiting BH_4_ synthesis in wild-type second-instar larvae by blocking the synthesis of melanin precursors in epidermal cells caused individuals to lose their original melanic body color (Lab unpublished contribution). The gene expressions were also significantly reduced when compared with that in the control group (Figure 5 bottom). Thus, the content and variation of the intracellular melanin precursors were important factors for regulating the expression of cuticular protein-encoding genes. We concluded that the inducing effect of the melanin precursors on the expression of cuticular protein-encoding genes is the basis for melanism promoting genes' transcription.

**Figure 5.**
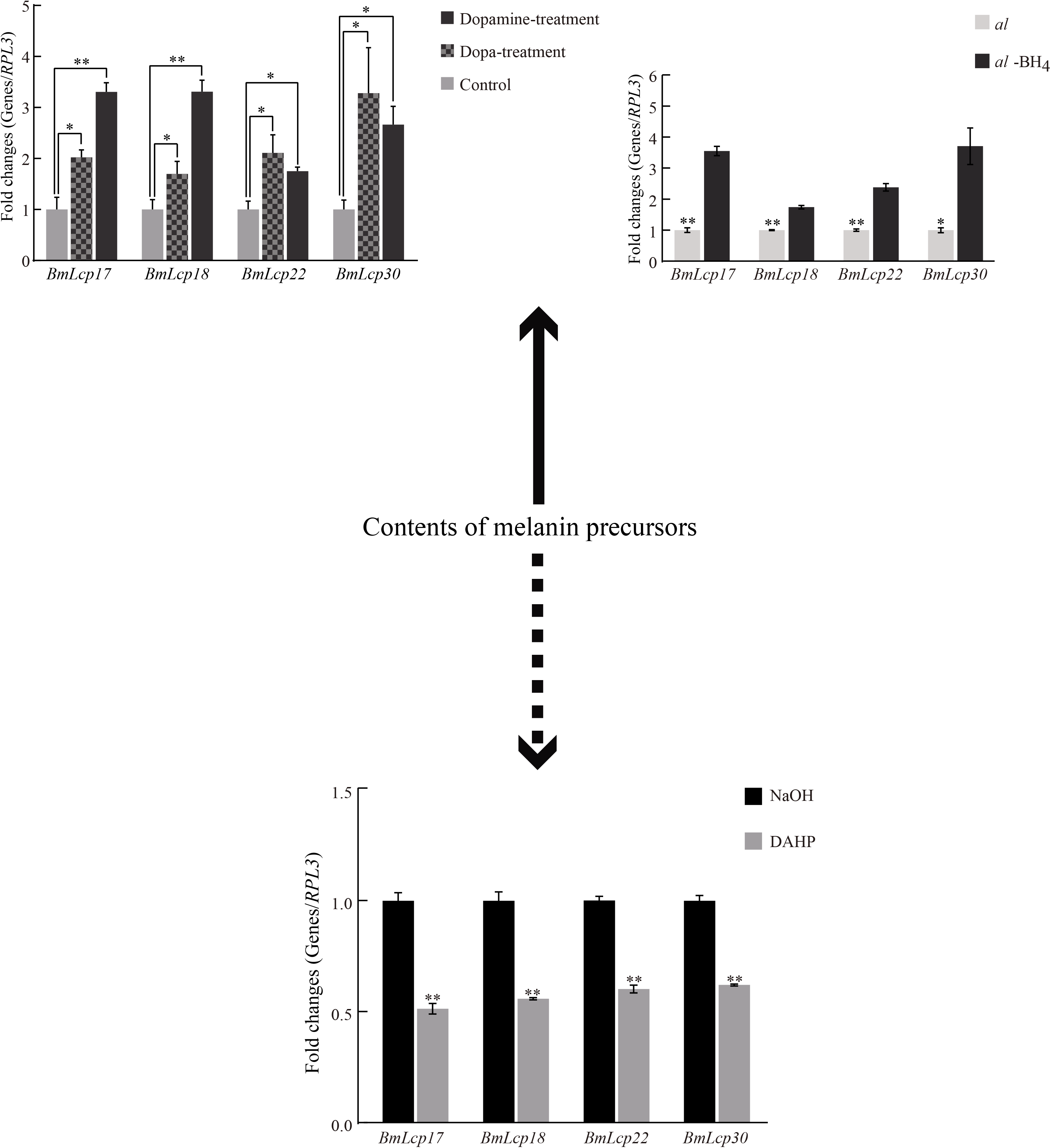
Effect of melanin precursors (top left: in cells) and BH_4_ (top right: in *vivo*) treatments on the expression of cuticular protein-enconding genes (*t*-test; n = 3, * *p* < 0.05; ** *p* < 0.01), and variations of gene expression levels in larval cuticle treated with the inhibitor DAHP (bottom). n = 3, *t*-test, ** *p* < 0.01.

## Discussion

Melanin is mainly deposited in the epicuticle and exocuticle, cross-linked with proteins in this layer, and involved in the formation of the melanic cuticular characteristics (Hopkins and Kramer 1992; Suderman *et al*. 2006; Andersen 2010; Moussian 2010; Hu *et al*. 2013). Some cuticular proteins located in the endocuticle may not be directly associated with the melanin, but the maintaining of the normal cuticle need to rely on the normal development and coordination between each cuticle layer, and these proteins are indispensable element in shaping the endocuticle (Moussian 2010; Cohen and Moussian 2016). Thus, we suppose that these endocuticular proteins may influence the transportation or accumulation of melanin through some indirect ways, to contribute to the unique features of the melanic cuticle. We indeed clearly observed that extensive star-like protrusions were present on the cuticle when a large amount of melanin accumulated (Figure S1). Similar correlations between cuticle structure and body color have been reported multiple times (Futahashi *et al*. 2012; Noh *et al*. 2016; Tan *et al*. 2016), imply that there should be some interactions between cuticular proteins and melanin. Although the exact interaction pattern between these two cuticular components is unknown, microscopic observation clearly shows that the deposition of an excessive amount of melanin affected the cuticle characteristics.

The expression profile of cuticular protein-encoding genes in melanic silkworm mutants and the black markings of *Papilio* larvae supported the hypothesis that up-regulated cuticular proteins probably participate in the transport or maintenance of the corresponding pigments in a specifically colored cuticles (Figures 1, 2 and Figure S1) (Futahashi *et al*. 2012; He *et al*. 2016; Wu *et al*. 2016). Yet over-expression of cuticular protein-encoding genes *in vivo* cannot trigger (or induce) the formation of melanic cuticle (Tajiri *et al*. 2017). Therefore, we hypothesized that if instructions for melanin accumulation were included in the developmental program for which melanism is the original factor, other cuticle features and structures would adapt to the level of melanin accumulation, regardless of genetic background. The up-regulation of cuticular protein-encoding genes should be a necessary adaptation for the maintenance and stability of the structural characteristics and physical properties of melanic cuticles. The relationship of melanin and cuticular proteins at the ultrastructural level deserves special attention in follow-up studies.

Although the elaborate regulatory mechanism by which melanin precursors affect the expression of cuticular protein-encoding genes is unclear, our results revealed the existence of this regulatory phenomenon (Figure 5). Cuticle formation is regulated stringently by temporal and spatial patterns, the accumulation and oxidization of melanin precursors, and interactions such as crosslinking among other components should occur after the initial cuticle formation (Moussian 2010; Sobala and Adler 2016; Tajiri 2017). Therefore, we propose that the cuticular proteins induced by melanin precursors are used to create a platform for further shaping and perfection of the melanic cuticle. When melanin-associated protein-encoding genes have similar expression patterns and functions (Nakato *et al*. 1994; Nakato *et al*. 1997; Shofuda *et al*. 1999; Okamoto *et al*. 2008; Liang *et al*. 2010; Tang *et al*. 2010; Qiao *et al*. 2014), the production of large amounts of functionally similar cuticular proteins would be driven by the melanic background to guarantee construction and stability of the melanic cuticle. During this process, melanism would unlocks the complementary features of melanin-associated cuticular proteins (such as *BmLcp18, BmLcp22, BmLcp30*), rescued cuticular malformation caused by the loss of function of some cuticular proteins (such as defected *BmLcp17* in *stony* mutant). Because a special cuticle-forming pattern appears to be regulated by melanin accumulation, melanism may enhance insect adaptability to avoid the impairment of survival caused by mutations and/or functional loss of some cuticular proteins, and our results also add new evidence to explain how melanism can be a beneficial trait (Wilson *et al*. 2001; True 2003; Wittkopp *et al*. 2003; Wittkopp and Beldade 2009; Liu *et al*. 2015; Mallet and Hoekstra 2016).

As far as we known, there is no evidence to suggest that the four larval cuticular proteins (as structure proteins) and their orthologous can enter the nucleus and regulate gene expression by acting as transcription factors. In addition, our findings and some other studies showed that changes in the expression of one cuticular protein-encoding genes of R&R family did not affect the expression of other members (Figure S6, S7) (Arakane *et al*. 2012; Noh *et al*. 2015). Moreover, several studies reveal that organisms optimize resources allocation at gene or protein expression level (Liebermeister *et al*. 2004; Dekel and Alon 2005), and there is no report that the cuticular proteins have long range morphogen effects (such as wingless) to regulate their encoding genes by feedback regulation from the cuticle to the epidermal cells; thus, the expression of melanin associated cuticle protein-encoding genes should be appropriately and simultaneously coordinated with the sufficient accumulation of intracellular melanin precursors, as a relatively direct, economical, efficient and convenient strategy. Furthermore, DAHP inhibits GTPCHI activity, but it does not directly impair the extracellular accumulation of melanin and protein–protein interactions in the cuticle. Finally, there is some evidence that melanin precursors regulate gene expressions via the receptors (Konradi *et al*. 1996; Berke *et al*. 1998; Westin *et al*. 2001). In summary, we hypothesized that the up-regulation of the four *BmLcp* genes were induced simultaneously by excessive amounts of melanin precursors, but should not be the interaction among these four genes on regulation level. To test our hypothesis, further analyses will be performed to determine the expression of the remaining *BmLcp* genes in some *BmLcps* mutant (such as defective *BmLcp17*) cell line by increasing or decreasing the melanin precursors content.

The markings were lighter in the *stony* mutant than in the wild-type strain, and were accompanied by the downregulation of melanin synthase genes (Figure S4). If the dysfunction of some cuticular proteins cannot be effectively supplemented, abnormalities of the cuticle structure and interactions of various cuticular components may occur, probably resulting in barriers to melanin deposition and metabolism. This dysfunction might be the reason why the *p^S^/+^pS^, st/st* (or *p^M^/+^pM^, st/st*) genotypes was lighter than the *p^S^*/+^*pS*^, +^*st*^/_ (or *p^M^*/+^*pM*^, +^*st*^/_) genotypes and had a different body shape. Because the homozygous of *p^B^* mutation is lethal (Xiang 1995; Banno *et al*. 2005), we were unable to obtain F_2_ progeny with the *p^B^*/*p^B^, st/st* genotype. However, we note that the *p^B^*/+^*pB*^, +^*st*^/*st* and *p^B^*/+^*pB*^, *st/st* genotype individuals had almost the same body shape and melasnism (Figure 4A). Therefore, we propose that during cuticle formation, if the signal promoting melanism is sufficiently strong, sufficient melanin precursors should be generated. The contents of functionally redundant proteins induced by melanin precursors are sufficient to fill the gap generated by the dysfunction of some cuticular proteins, guaranteeing the normal accumulation of melanin. Therefore, we hypothesize a threshold for the melanism-promoting effect. When the accumulation of melanin precursors spans this threshold, the requirements for cuticular proteins with similar function are not lowered, even when one of them loses the function. This effect would guarantee the formation of a normal cuticle structure, ensuring subsequent pigmentation.

We propose the following regulatory model: when large amounts of melanin precursors induced by endogenous- and/or exogenous melanism-promoting factors, it may directly or indirectly induce the up-regulation of some cuticular protein-encoding genes. This upregulation guarantees the formation of normal structural features of the melanic cuticle. During this process, if some cuticular proteins lose function, other functionally redundant cuticular proteins induced by melanin precursors compensate for the functional defects. Compensatory intensity increases with increasing melanin accumulation. When melanin accumulation spans a certain threshold, compensation completely masks the defective phenotype caused by the malfunctioning genes. This model adds to growing evidence that melanism may have pleiotropic effects that enhance adaptability over and above the effect of melanin accumulation itself (Figure 6). Due to the coexistence of excess melanin and cuticular proteins is common in other insects, and homologues the four *BmLcp* gene are widely distributed in the Lepidoptera (Table S2), we presume that the above reciprocal action and its corresponding biological significance are conserved in other Lepidoptera insects to that in *Bombyx mori*.

**Figure 6.**
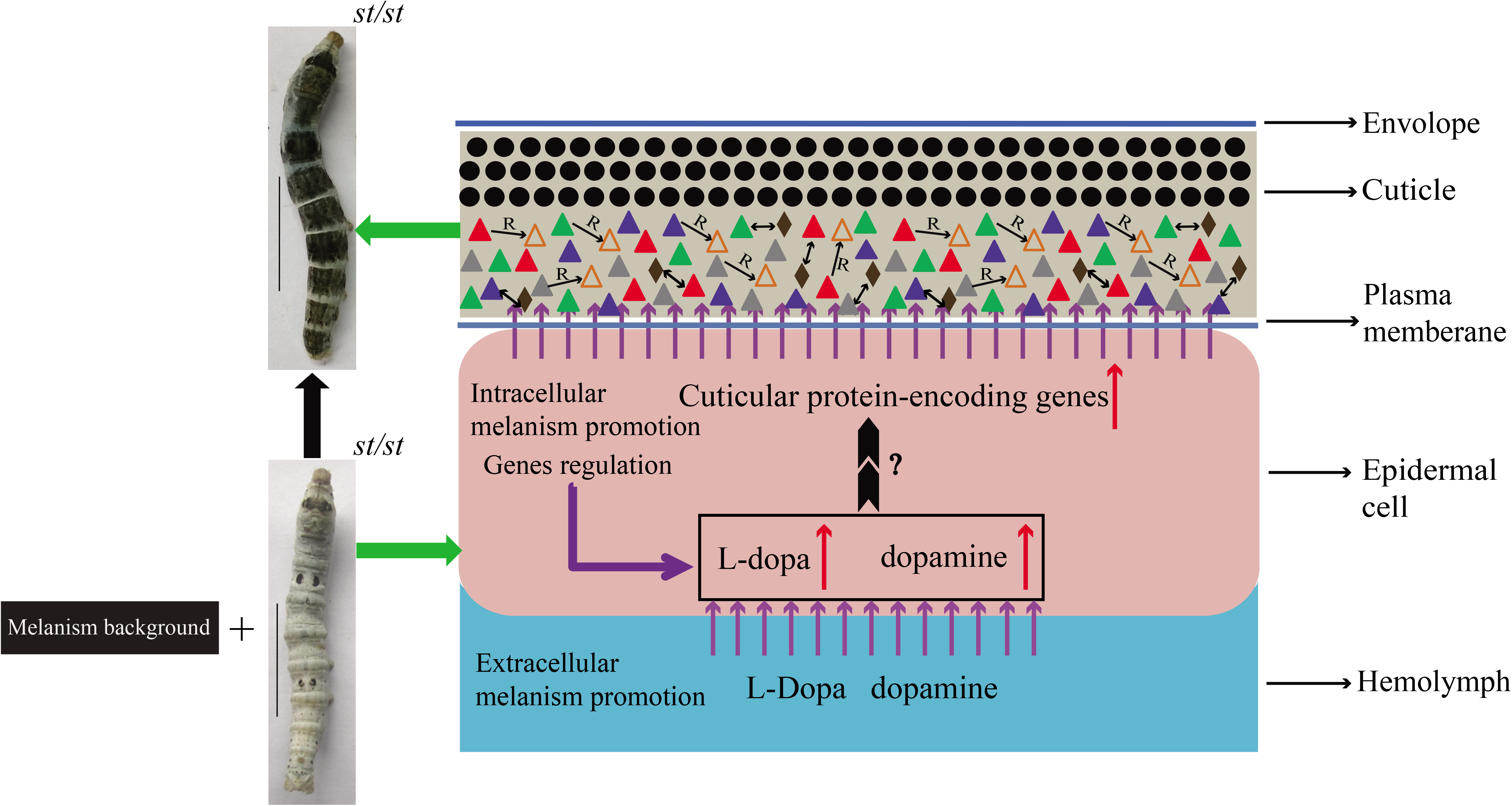
Schematic overview of the effect of melanin precursors on the expressions of cuticular protein-encoding genes. Black solid circles represent the melanin. Triangles represent the BmLcps with similar expression patterns and functions. Solid and hollow triangles represent the normal and defective functions, respectively. Brown rhombi represent other components (such as chitin) in the cuticle. Solid double-headed arrow indicates the probable interaction between the endocuticular proteins and other components. Combination of the single-headed arrow and the letter ‘R’ indicate the repair of the potential defects through functional complementary. Purple arrows show the direction in which the melanin precursors or cuticular proteins flow from the haemolymph to the epidermal cells as well as from the epidermal cells to the cuticle. Red arrows indicate the increased contents of melanin precursors or the up-regulation of cuticular protein-encoding genes. Purple polyline arrows indicate melanism-promoting factors produced by other genetic instructions. Double dovetail arrows indicate the effect of melanin precursors on cuticular protein-encoding genes. The question mark indicates that the details of induction (directly or indirectly) are unclear.

In summary, we used crucial cuticle components to elucidate the mutual effects among some key genetic factors, and the physiological significance of these mutual effects during the development of the morphological features. Our findings contribute a realistic basis for in-depth study on the interaction patterns of melanin and cuticular proteins, and will also guide relevant studies in other Lepidopteran insects or other insect species.

## Acknowledgments

We thank Dr. Tianzhu Xiong, Dr. Yonggang Hu and Ms. Yan Yao for the valuable advices. This work was supported by the Hi-Tech Research and Development 863 Program of China (Grant No. 2013AA102507), the National Natural Science Foundation of China (No. 31302038; 31372379), the Natural Science Foundation Project of ChongQing (CSTC) (No. cstc2013jcyjA80022), China Scholarship Council (201508505020) and Par-Eu Scholars Program (20136666).

